# Lipolytic and Anti-Adipogenic Effect of Capsaicin, Camphor, and Caffeic acid on Human SGBS Adipocytes

**DOI:** 10.1101/2025.05.05.652232

**Authors:** Uzair Ali, Martin Wabitsch, Danijela Gregurec

## Abstract

Excess fat accumulation contributes to metabolic disorders such as insulin resistance, type 2 diabetes, cardiovascular disease and increases risk of dementia. Promoting lipolysis and inhibiting adipogenesis through natural compounds offers a promising therapeutic approach to obesity. This study investigates the lipolytic and anti-adipogenic effects of capsaicin, camphor, and caffeic acid in human Simpson-Golabi-Behmel Syndrome (SGBS) adipocyte model. First, we determined appropriate treatment concentrations using MTT assays, which demonstrated a dose dependent reduction in cell viability for all three compounds. Selected doses were applied to differentiating SGBS cells until day 14. Lipid droplet accumulation and free fatty acid release were assessed using Oil Red O (ORO) staining and a lipolysis assay kit, respectively. Gene expression of adipogenic and lipolytic markers was analysed by RT-PCR and TRPV1 receptor involvement was examined by immunofluorescence. A network pharmacology approach incorporating GeneCards, STITCH, and other databases revealed downregulation of PPARG, CEBPA, and FABP4, and upregulation of HSL, ATGL, and PLIN1. TRPV1 activation was prominent in capsaicin treated cells. Network analysis identified shared regulatory hubs such as PPARG, STAT3, and MTOR. All treatments significantly reduced lipid accumulation and increased lipolysis, with capsaicin showing the strongest effects. Combination treatments, especially capsaicin with caffeic acid, exhibited synergistic effects. While the previous studies showed interference of these compounds in the molecular pathways involved in fat cell metabolism, our work establishes for the first time potential thermogenic capacity of camphor individually and of combinatory effects of three compounds to inhibit adipogenesis and promote lipolysis in human adipocytes, potentially through TRPV1 signalling and modulation of metabolic gene networks. These findings highlight their therapeutic potential for metabolic disorders.

## 1. Introduction

Obesity is a complex metabolic disorder characterized by excessive fat accumulation, primarily because of an imbalance between energy intake and expenditure^1^. The inflammation and pathogenesis of obesity is closely related to adipose tissue expansion which mainly occurs through hypertrophy (enlargement of adipocytes), and hyperplasia (increase in adipocytes population)^2^. Targeting adipose tissue metabolism by promoting lipolysis and regulating the adipogenesis has emerged as potential strategy for mitigating obesity and its associated disorders^3,4^. For instance, obesity is a major risk factor for metabolic syndrome, which increases susceptibility to developing chronic conditions including type 2 diabetes, hypertension and cardiovascular diseases^5^. It has also been increasingly associated with neurological disorders, including cognitive decline, Alzheimer’s disease, and cerebrovascular diseases^6^. Studies suggest that obesity linked metabolic dysfunction contributes to insulin resistance, vascular dysfunction and neuroinflammation which aggravate cognitive impairment and increase the risk of dementia^7, 8^.

Natural bioactive compounds have gained considerable attention for their ability to modulate fat loss through interference in metabolic pathways and their relatively low toxicity^9^. Among these compounds that can regulate adipocyte function are camphor, capsaicin, and caffeine^10^. Studies indicate that camphor influence mitochondrial activity, intracellular cAMP level, and increase thermogenesis in adipose tissue, thereby regulating metabolic activity and increasing energy expenditure^11^. Capsaicin, an active alkaloid compound found in chilli pepper (capsicum annuum), is a known agonist for transient receptor potential cation channel subfamily V member 1 (TRPV1) which plays a vital role in regulating energy metabolism and lipid oxidation through the activation of sympathetic nervous system^12, 13, 14^. Capsaicin has attracted significant interest in fat metabolism given its ability to activate TRPV1 and increase energy expenditure^15^. Caffeine, a methylxanthine has been reported to promote the breakdown of triglyceride and the release of free fatty acid by activating PKA and HSL through increased intracellular cAMP level in 3T3-F422A adipocyte model^16, 17^.

While the effect of these compounds has been shown to modulate fat metabolism in various cells and animal models, their comparative effects in human adipocytes models remain to be understood. There are few human adipocyte cell models available, with SGBS cells being example with prolonged proliferation and differentiation capacity^18^. Unlike other immortalized cell lines, SGBS cells can differentiate into metabolically active mature adipocytes with similar characteristics to human derived primary adipocytes^19^. Given the physiological relevance of SGBS cells as an *in vitro* model of human adipocyte model, investigating bioactive compounds can provide important insight into their regulatory effects on fat metabolism^20, 21^. In this study, SGBS cells are employed as a human adipocyte model to evaluate bioactive compounds for their ability to promote lipolysis and inhibit adipogenesis. In addition to capsaicin and caffeic acid, we used camphor and the combination treatments of all three compounds for the first time to evaluate their effect on promoting lipolysis and inhibiting adipogenesis.

## 2. Materials and Methods

### 2.1 Reagents

Reagents were obtained from Sigma Aldrich or Thermo Fisher Scientific, unless otherwise stated. Rosiglitazone was purchased from Cayman Chemical. The LIVE/DEAD^TM^ Cell Imaging Kit was from Abcam. Primers were synthesized by Eurofins Genomics. Anti TRPV1 primary antibodies and CFL647-conjugated secondary antibodies were obtained from Santa Cruz Biotechnology.

### 2.2 Cell culture

Human SGBS cells were kindly provided by prof. M. Wabitsch (University of Ulm, Germany). Cells were grown in dulbecco modified eagle’s medium F-12 (DMEM/F-12) supplemented with 10% fetal bovine serum (FBS), 3.3 mM biotin, 1.7 mM pantothenate and 1% penicillin/streptomycin solution, at 37°C with 5% CO_2_ and 95% relative humidity. Differentiation was induced on 70% confluent cells with serum free growth medium supplemented with 10 µg/ml transferrin, 0.2 nM triiodothyronine (T_3_), 250 nM hydroxycortisone, 20 nM human insulin, 25 nM dexamethasone, 250 µM 3-isobutyl-1-methylxanthine (IBMX) and 2 µM rosiglitazone (day 0 of differentiation). After 4 days, the differentiation medium was replaced with a maintenance medium that lacking IBMX, dexamethasone and rosiglitazone. Fresh maintenance medium was added every 4 days.

### 2.2 MTT assay

Cell Viability MTT assay was performed to determine a nontoxic dose for cells treatment. Compounds were diluted in serum free DMEM/F12 and added for 96 h. MTT (20 µl, 5 mg/ml) was added for 4 h, then media removed, and formazan crystals dissolved in DMSO. Absorbance at 570 nm was measured with Spectramax M2 reader to assess viability.

### 2.4 LIVE/DEAD assay

Live/Dead assay was performed on differentiated mature adipocytes following treatment with selected concentrations from day six until day 14 of differentiation. The treatments groups included capsaicin (CAP, 1µM), camphor (CAM, 0.1Mm), and caffeic acid (CA, 1mM) as well as their combinations (CAP-CAM (1µM – 0.1mM), CAM-CA (0.1mM – 1mM), and CAP-CA (1µM – 1mM). Untreated cells with only ethanol in the media served as a control. On day 14, the assay was performed using the LIVE/DEAD assay kit (Abcam), following the manufacturer’s instructions. Images were taken with Olympus IX 73 inverted fluorescence microscope with cellSens imaging software using FITC (excitation 495 nm, emission 519 nm) and Texas Red excitation 595 nm, emission 615 nm) filters.

### 2.5 Adipogenesis assay

To assess the anti adipogenic effect of capsaicin, camphor, and caffeic acid, at day 6 of differentiation, treatments were initiated and continued until day 14. Treatments were selected based on the MTT assay results by adding it to the maintenance media at day 6 of differentiation which was refreshed after every 2-3 days until day 14. Then cells were fixed in 4% PFA, dehydrated with 60% isopropanol, and stained with Oil Red O (3:2 in water) for 20 minutes. After washing with distilled water, images were taken using an inverted microscope. For the quantification of lipid contents, ORO was extracted with 100% isopropanol and absorbance was measured at 490nm using Spectramax M2 microplate reader.

### 2.6 Lipolysis assay

To examine whether capsaicin, camphor and caffeic acid can induce lipolysis in differentiated SGBS cells, the cells were treated with chosen concentrations at day 12 until day 14. Positive control was treated with 10μM isoproterenol for 3 hours on day 14 to induce maximum release of free fatty acids. Lipolysis was measured using the Abcam lipolysis assay kit (Abcam, USA) following the manufacturer’s protocol. A standard curve was plotted by graphing absorbances against the known glycerol concentration from the standards. This produced a linear aggression equation which for determining glycerol concentrations.

### 2.7 Gene expression

Total RNA was extracted using TRIzol**^TM^** Reagent (Thermo Fisher Scientific, USA) following the manufacturer’s protocol. RNA concentration and purity were assessed using a NanoDrop**^TM^** 2000 spectrophotometer (Thermo Fisher Scientific, USA). Primers were designed using NCBI Primer BLAST tool (**Tab. S1**). One microgram of total RNA per sample was reverse transcribed into complementary DNA (cDNA) using the High Capacity cDNA Reverse Transcription Kit, following the manufacturer’s protocol. Quantitative real time PCR (qPCR) was conducted using the qTOWER³ real time PCR system (Analytik Jena, Germany) with SYBR Green master mix. All primers were pre validated for amplification efficiency (range 90 – 110%) before use in expression analysis. Target genes included markers of adipogenesis such as PPARγ, ADIPOQ, FABP4, CEBPα and key regulators of lipolysis such as HSL, PLIN1, and ATGL/PNPLA2 were quantified for expression in response to treatments. Peroxisome proliferator activated receptor gamma coactivator 1-α (PGC1α) was quantified for metabolic and mitochondrial activity during adipogenesis and lipolysis. Gene expression was normalized using housekeeping genes β-actin (ACTB) and RPLP0. Relative gene expression was calculated using the 2^−ΔΔCt method to calculate fold change versus control group.

### 2.8 Immunofluorescence staining

To investigate the effect of selected bioactive compounds on TRPV1 expression during adipogenesis and lipolysis, cells were treated with capsaicin, camphor, and caffeic acid at the same concentration and treatment duration used for adipogenesis and lipolysis. Untreated differentiated cells served as controls. At day 14, cells were fixed in 4% paraformaldehyde, permeabilized with 0.1% Triton X-100, and blocked with 3% normal serum in PBS at room temperature to prevent nonspecific antibody binding. TRPV1 expression was evaluated via immunofluorescence using a TRPV1 primary antibody (1:200), and Alexa Fluor 647 conjugated secondary antibody (1:500). Lipid droplets were stained using BODIPY 493/503 (1 µg/mL) and nuclei with DAPI. Coverslips were mounted using antifade mounting medium and images were capture with Olympus IX73 inverted fluorescence microscope. TRPV1 expression and lipid droplet morphology were assessed qualitatively based on fluorescence signal intensity and distribution patterns.

### 2.9 Target gene predictions

To predict target genes for capsaicin, camphor, and caffeic acid, four online tools PharmMapper, SEA, STITCH, and Swiss Target Prediction were used. Compound structures and SMILES were obtained from PubChem and converted to .sdf format for PharmMapper. Predicted targets were standardized via UniProt (Homo sapiens). Genes related to lipolysis, adipogenesis, and thermogenesis were retrieved from GeneCards. Overlapping genes were identified using a Venn diagram tool, and functional networks were constructed in Cytoscape using GeneMANIA. Network topological analysis was performed using the CytoNCA plugin in Cytoscape, with centrality parameters including Degree Centrality (DC), Betweenness Centrality (BC), and Closeness Centrality (CC), all calculated using the “without weight” option. The top 10 hub genes were identified based on these centrality scores which were further evaluated using VarElect to explore their relevance to lipolysis, adipogenesis, and thermogenesis based on -log₁₀(p) and disease likelihood scores.

### 2.10 Statistical analysis

All statistical analyses were performed using XLSTAT software. For gene expression, unpaired t-test were performed to compare each treatment with the control for every gene, and p value were adjusted using the Benjamini Hochberg False discovery rate (FDR) method to account for multiple comparisons. Data from qRT-PCR, cell viability, lipolysis, and adipogenesis assays are presented as mean ± standard error of the mean (SEM). Statistical significance was defined as p < 0.05 (), p < 0.01 (), and p < 0.001. Data were visualized using bar graphs with error bars, with asterisks indicating statistically significant differences. For all experiments, biological replicates were performed in triplicate (n = 3 per condition). For qRT-PCR, each biological replicate was analyzed in three technical replicates (n = 9 per condition).

## 3. Results

### 3.1. Cell Viability assays

The MTT assay showed a concentration dependent decrease in SGBS cell viability after 4 days of treatment with capsaicin, caffeic acid, and camphor (**Fig. 2a–c**). Capsaicin maintained high viability at 1 µM (94.17%) and 5 µM (85.11%) but showed strong cytotoxicity at 50 µM (21.17%, p < 0.001). Caffeic acid reduced viability to 90.73% at 0.1 mM and 86.80% at 1 mM, with significant drops at 5 mM (53.56%, p < 0.01) and 10 mM (21.78%, p < 0.001). Camphor caused a moderate viability reduction, with 88.48% at 0.01 mM and 72.85% at 1 mM (p < 0.001). Based on these results, a suitable concentration of each compound was chosen to examine their anti adipogenic and lipolytic effect in the follow up experiments. However, to further confirm the suitability of the selected concentrations for differentiating SGBS cells from day six to day fourteen of differentiation, a LIVE/DEAD assay was performed, based on calcein staining of live cells and ethidium homodimer-1 (EthD-1) staining of dead cells (**Fig. 2e**). The results showed no significant difference in cell viability among the treated and untreated cells indicating the suitability of the selected concentrations for further experiments (**Fig. 2d**).

**Figure 1.**
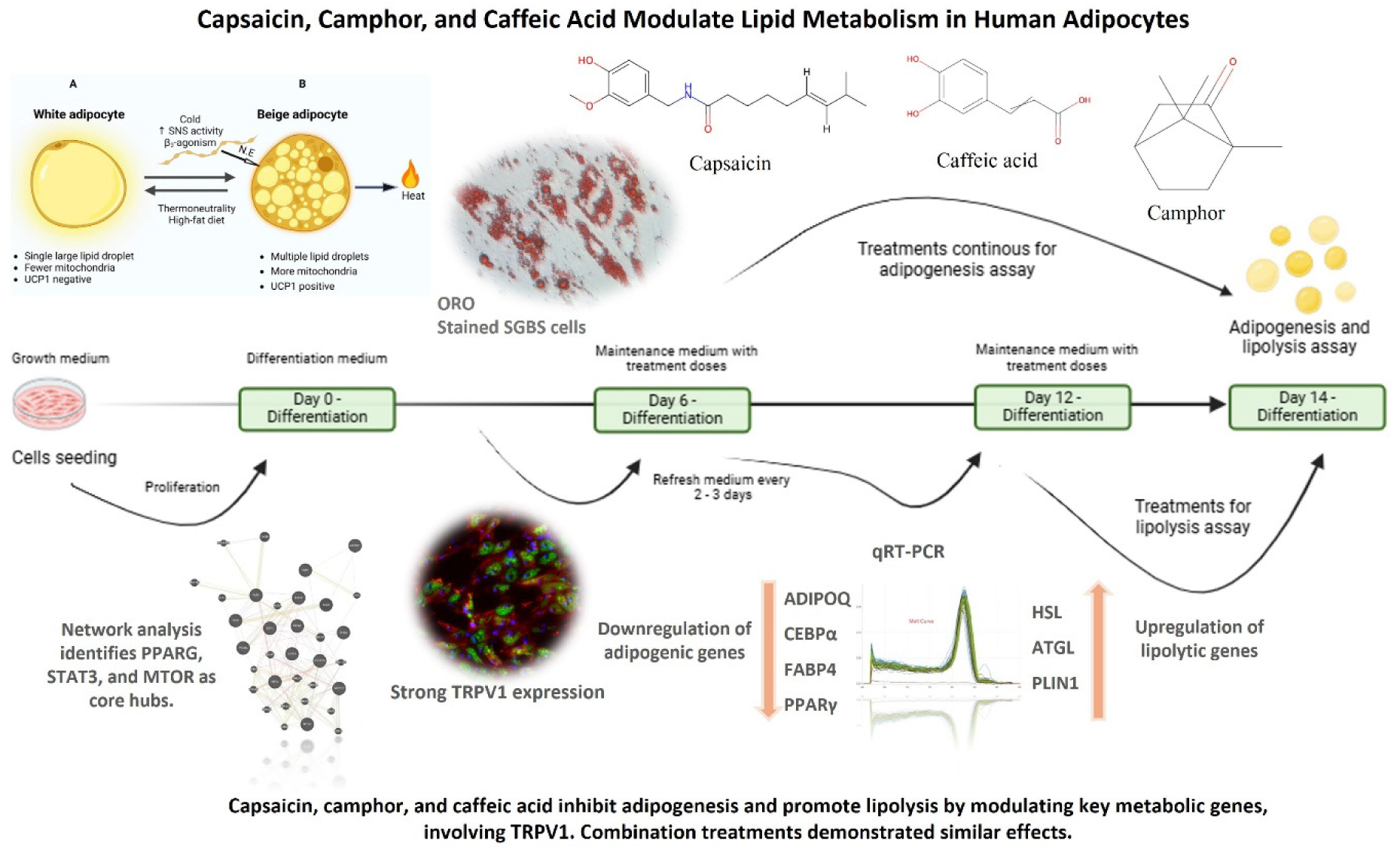
Graphical overview of the experimental design and findings. Differentiating SGBS cells were treated with capsaicin, camphor, and caffeic acid, and their combination, to assess lipolytic (Day 12–14) and anti adipogenic effects (Day 6–14). Treatments reduced lipid accumulation, enhanced lipolysis, and modulated the expression of key metabolic genes. Network analysis revealed PPARG, STAT3, and MTOR as central regulatory hubs, while immunofluorescence indicated TRPV1 involvement, particularly with capsaicin. Combination treatments exhibited comparable effects.

**Figure 2.**
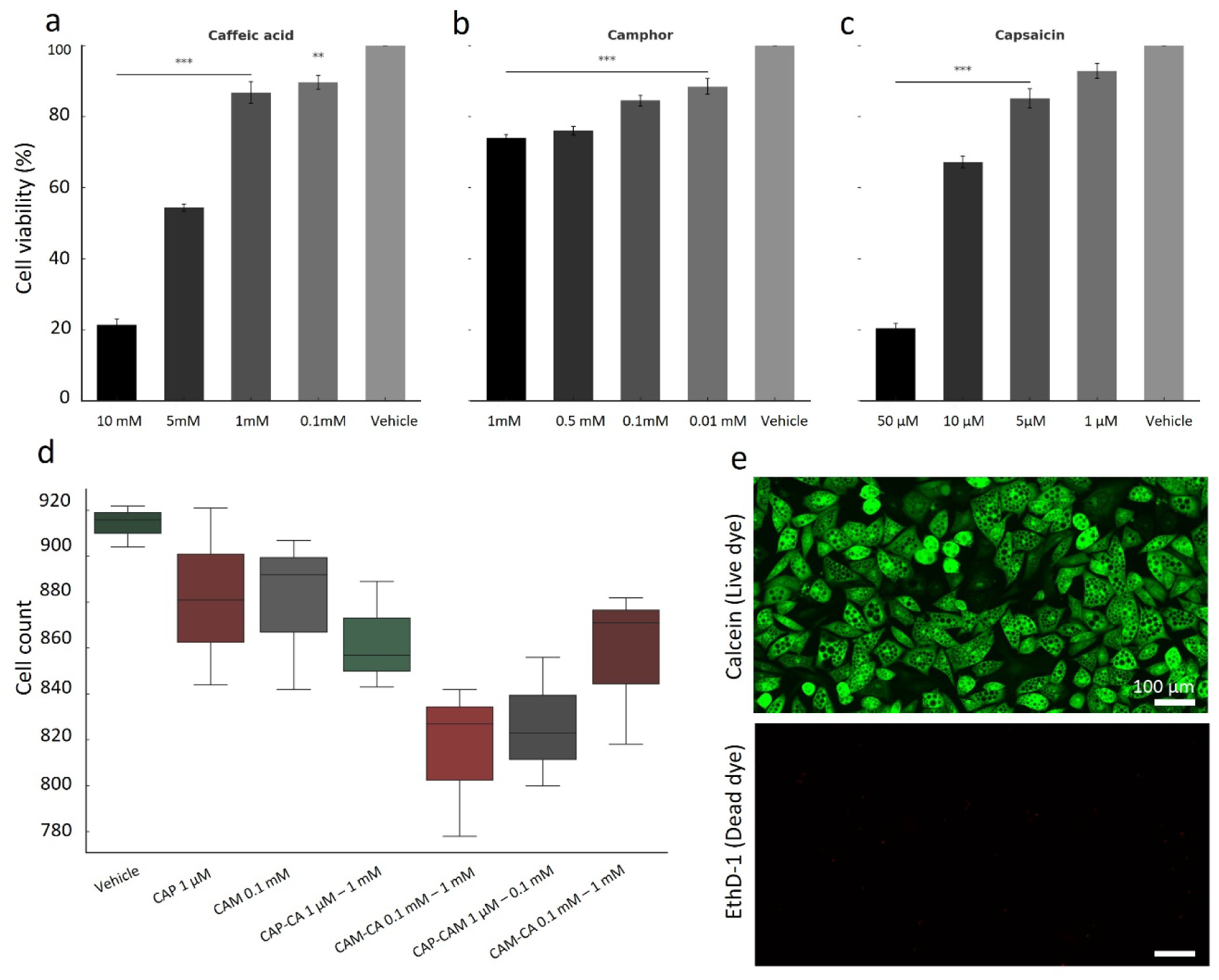
Effects of caffeic acid, camphor, and capsaicin on cells viability. **(a – c)** MTT assay results showing dose dependent effects of **(a)** caffeic acid, **(b)** camphor, and capsaicin **(c).** High concentrations of capsaicin (50 μM), camphor (1 mM), and caffeic acid (10 mM) significantly reduced cell viability (P < 0.001), while lower doses showed minimal cytotoxicity and maintained viability comparable to vehicle. **(d)** Live cell quantification from Calcein-AM staining (LIVE/DEAD assay). No significant effect on live cell counts. Statistical analysis was performed using one-way ANOVA followed by Tukey’s post hoc test. **(e)** Representative fluorescent micrograph of Calcein-stained live cells (green), confirming qualitative viability consistent with quantitative findings in panel (d). Scale bar = 100 μm.

### 3.2 Lipid accumulation in SGBS cells reduced in response to the treatments

Microscopic images showed that treatment with capsaicin, camphor, and caffeic acid resulted in reduced lipid droplet formation compared to untreated control cells differentiated under normal physiological conditions (**Fig 3a**). A significant decline in lipid content accumulation from the baseline 100% vehicle treated control cells was observed (**Fig 3b**). Among the treatments, capsaicin at 1 µM concentration induced the most significant effect reducing lipid accumulation to 50.67% (p < 0,001). Camphor (0.1 mM) and caffeic acid (1 mM) were moderately effective, lowering lipid droplets formation to 64.08% and 68.91% respectively (p < 0.01). Among the combination treatments, capsaicin and camphor (1 µM, 0.1 mM) inhibited lipid accumulation to 53.82% (p < 0.01%), while camphor and caffeic acid (0.1 mM, 1 mM) and capsaicin and caffeic acid (1 µM, 1 mM) resulted in 66.29% and 54.60% respectively. Capsaicin in combination with caffeic acid exhibited a comparable effect to capsaicin alone showing a high significance (p < 0.001), whereas combination of camphor and caffeic acid remained significant but less potent (p < 0.01) (**Fig 3b**).

**Figure 3.**
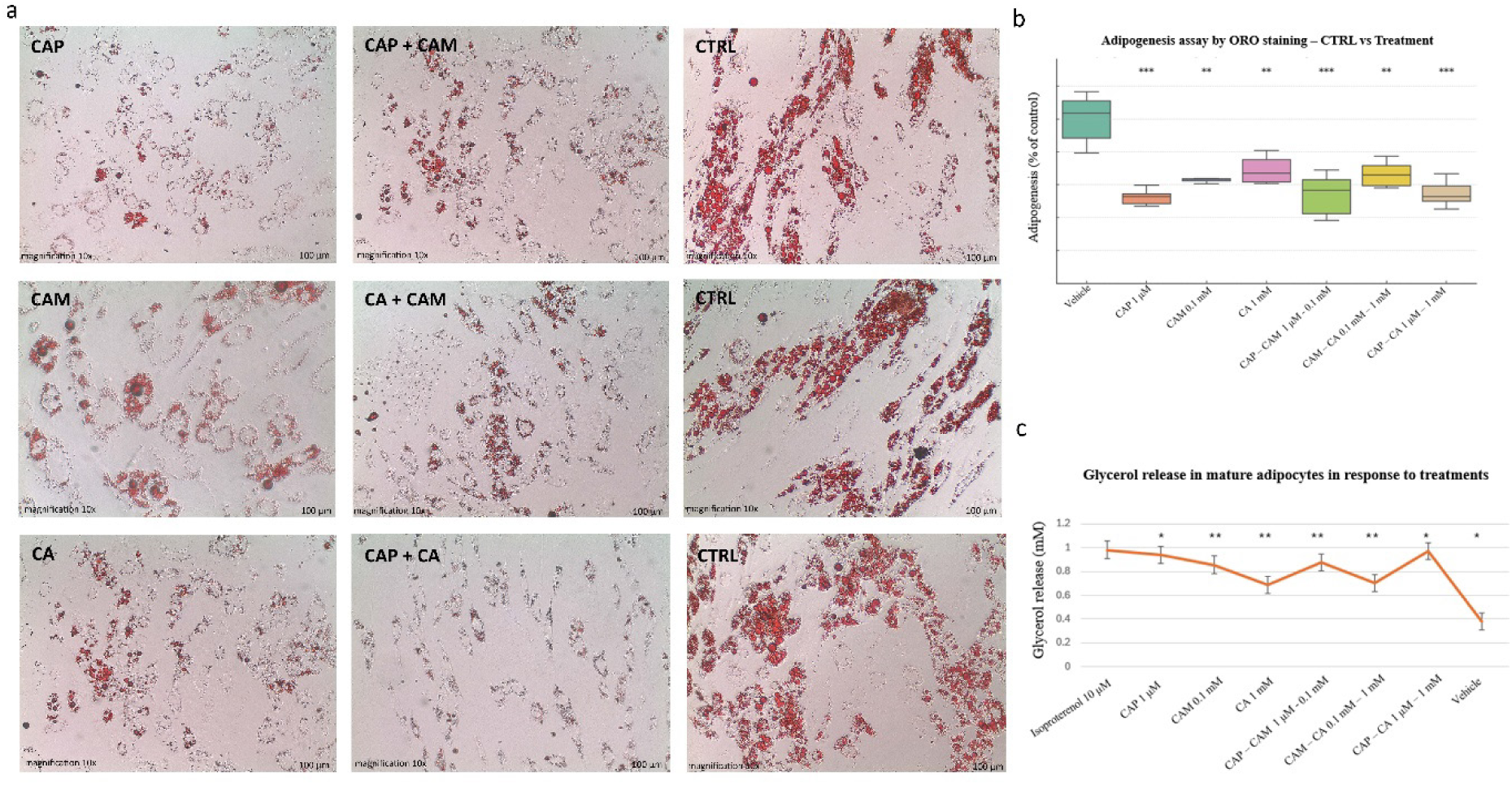
Effects of capsaicin, camphor, and caffeic acid on adipogenesis and lipolysis in SGBS cells. **(a)** Oil Red O (ORO) staining of differentiated SGBS cells treated with capsaicin (1 μM), camphor (0.1 mM), and caffeic acid (1 mM), individually and in combination. All treatments reduced lipid accumulation compared to control. Scale bar = 100 μm. **b)** Quantification of adipogenesis (% of control) based on ORO absorbance. All treatments significantly decreased adipogenesis compared to vehicle (*P* < 0.01 to *P* < 0.001; one-way ANOVA with Tukey’s post hoc test). **c)** Glycerol release was significantly increased by individual and combined treatments, with capsaicin and its combinations showing the strongest effects, comparable to the β - adrenergic agonist isoproterenol (*P* < 0.05 to *P* < 0.01).

### 3.3 Capsaicin, camphor, and caffeic acid promote lipolysis

Glycerol release is commonly used as a marker of lipolysis in lipid metabolism research. Our results of the adipolysis assay demonstrate that capsaicin at 1 µM, camphor 0.1 mM, caffeic acid 1 mM, and their combinations significantly influence glycerol release in differentiated SGBS adipocytes after 48 hours of treatment (**Fig 3c**). Isoproterenol a known β-adrenergic agonist at 10 µM concentration resulted in the highest glycerol release (0.98 mM, p < 0.05) compared to untreated control cells (0.38 mM). Among the tested compound, capsaicin significantly stimulated glycerol release (0.94 mM, p < 0.05) demonstrating receptor mediated lipolysis in mature SGBS cells. Similarly, camphor significantly stimulated glycerol release (0.86 mM, p < 0.05), The effect of caffeic acid was mild but significantly promoted lipolysis (0.69 mM, p < 0.05), Nonetheless, capsaicin in combination with caffeic acid and camphor demonstrated to promote lipolysis significantly higher (0.97 mM, p < 0.01, 0.88 mM, p < 0.01 respectively) and nearly equivalent to the positive control isoproterenol treatment. This indicates a synergistic interaction between capsaicin and other compounds leading to increases fat breakdown potentially through the activation of lipolytic pathways. In summary, the results suggest that capsaicin, camphor, and caffeic acid individually and in combination significantly enhance glycerol release, with CAP-CA demonstrating the strongest effect.

### 3.4 Effects of Capsaicin, Camphor, and Caffeic Acid on Adipogenic Gene Expression

Our data demonstrated that capsaicin induced the strong suppression of adipogenic markers. Expression of FABP4 was reduced to 0.22-fold (p < 0.001), while PPARG, ADIPOQ, and CEBPA were downregulated to 0.42-fold (p < 0.001), 0.41-fold (p < 0.001), and 0.25-fold (p < 0.001), respectively. These results represent the strongest transcriptional inhibition observed in the study (**Fig. 4b**). Camphor treatment resulted in a moderate yet consistent reduction in gene expression. PPARG and FABP4 levels decreased to 0.58-fold (p < 0.001) and 0.35-fold (p < 0.001), respectively, while ADIPOQ and CEBPA were reduced to 0.66 fold (p < 0.001) and 0.52-fold (p < 0.001), respectively. Similarly, Caffeic acid exhibited a suppression profile like camphor, though with slightly stronger inhibition of FABP4 at 0.23-fold (p < 0.001). PPARG and ADIPOQ levels declined to 0.43-fold (p < 0.001) and 0.70-fold (p < 0.001), respectively. Combination treatments potentiated the inhibitory effects of individual compounds. The CAP-CA and CAP-CAM groups displayed further reductions in adipogenic gene expression compared to camphor or caffeic acid alone. The CAM-CA combination was comparatively less potent but still significantly downregulated all four markers, with fold changes ranging from 0.39 to 0.48 (p < 0.001) (**Fig. 4b**). In summary, all treatments significantly suppressed adipogenic gene expression, with capsaicin demonstrating the most notable and consistent downregulation across all genes assessed.

**Figure 4.**
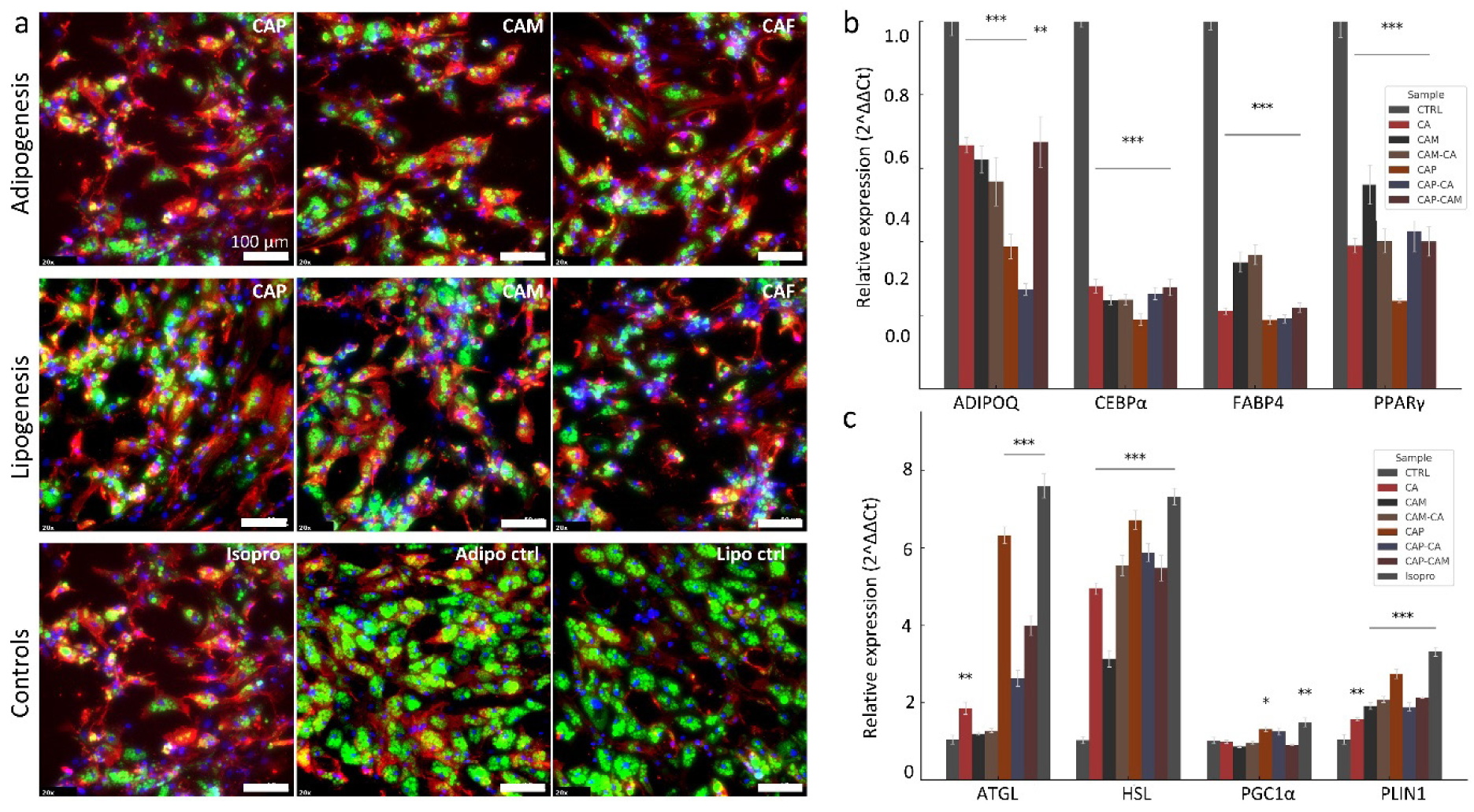
Effects of capsaicin, camphor, and caffeic acid on genes relevant to adipogenesis, lipolysis and intracellular TRPV1 expression in SGBS adipocytes. Immunofluorescence staining (**a**) and RT-qPCR analysis (**b, c**) of SGBS adipocytes treated with capsaicin (CAP), camphor (CAM), caffeic acid (CA), and their combinations from day 6 to day 14 of differentiation (adipogenesis phase) or day 12 to day 14 (lipolysis phase). RT-qPCR analysis of adipogenic genes (ADIPOQ, CEBPα, FABP4, PPARγ) showed that all treatments significantly downregulated expression, with CAP and its combinations (CAP-CA, CAP-CAM) producing the strongest inhibition (*P* < 0.001). **(c)**. Lipolytic gene expression (ATGL, HSL, PLIN1, PGC1α) was significantly upregulated by all treatments, particularly CAP and its combinations, with levels approaching those seen with the isoproterenol (Isopro) positive control. Data are presented as mean ± SEM (n = 9); *P* < 0.05, *P* < 0.01, *P* < 0.001.

### 3.5 Capsaicin, Camphor, and Caffeic Acid Stimulate Lipolytic Gene Expression

All treatments significantly upregulated lipolytic genes, though the extent of induction varied across conditions. Capsaicin produced the most substantial activation among single compound treatments (**Fig. 4c**). HSL expression increased to 6.73-fold, ATGL to 6.52-fold, and PLIN1 to 2.79-fold (all p < 0.001). PGC1A was also modestly induced, reaching 1.30-fold (p < 0.05), suggesting potential activation of mitochondrial biogenesis. Camphor showed a more modest effect, with HSL, ATGL, PLIN1 levels rising to 3.14-fold, 1.42-fold, and 2.00 fold respectively (both p < 0.001). whereas PGC1A expression largely unchanged. Caffeic acid demonstrated greater responses than camphor but was less effective than capsaicin. HSL increased to 4.95-fold (p < 0.001), ATGL to 2.00-fold (p < 0.01), and PLIN1 to 1.69-fold (p < 0.01). A moderate increase in PGC1A to 1.28-fold was also observed (p < 0.05).

Combination treatments showed similar pattern of enhanced gene expression with HSL increasing to ∼5.4–5.9-fold and ATGL to ∼3.2–4.3-fold across CAP-CAM, CAP-CA, and CAM-CA groups (all p < 0.0001). PLIN1 expression rose ∼2.1–2.2-fold, while PGC1A showed modest increases (∼1.3-fold; p < 0.05). As expected, isoproterenol induced the highest expression levels, with HSL and ATGL exceeding 7-fold. In summary, all tested compounds significantly activated lipolytic gene expression.

### 3.6 High TRPV1 expression in response to treatments

Immunofluorescence revealed that TRPV1 was expressed in both control and treated SGBS cells, with markedly enhanced signal intensity observed in response to the bioactive compounds’ treatments (**Fig 4a**). In the adipogenesis assay, capsaicin treated cells exhibited the strongest TRPV1 fluorescence, suggesting strong activation of the ion channel. Expectedly, camphor treatments also induced increased TRPV1 expression, but interestingly, caffeic acid treated cells showed similar strong signal compared to the untreated control, although the intensity was less pronounced than with capsaicin. Similarly, during the lipolysis phase (**Fig. 4a**), capsaicin treated cells again showed the most prominent TRPV1 signal, followed by camphor and caffeic acid, relative to the control. This pattern of elevated TRPV1 expression across both adipogenesis and lipolysis duration supports the hypothesis that TRPV1 may be a critical mediator of the observed bioactive effects.

Concurrently, BODIPY staining (green) demonstrated clear differences in lipid droplet morphology between control and treated cells (**Fig. 4a**). In the adipogenesis treatment groups, a reduction in the size and abundance of lipid droplets was observed, with many cells exhibiting numerous small, dispersed lipid droplets instead of the large, unilocular droplets characteristic of mature adipocytes in control samples. During the lipolysis treatment phase, a similar decrease in lipid droplet size and content was evident, supporting the notion of active lipid breakdown.

### 3.7 Identification and Functional Characterization of Hub Genes in Adipocyte Metabolic Networks

Network pharmacology analysis combined with topological modelling revealed central gene regulators potentially mediating the metabolic effects of capsaicin, camphor, and caffeic acid. Genes associated with adipogenesis (n = 1789), lipolysis (n = 816), and thermogenesis (n = 600) were initially retrieved from GeneCards, and their intersection yielded 150 common genes involved in all three processes (**Fig. 5d**). These 150 genes were subsequently intersected with predicted molecular targets of each compound, as identified through STITCH, Swiss Target Prediction, and SEA databases. Across all three compounds, several genes emerged as consistently central to the network structure. STAT3, ESR1, and MTOR were prominently ranked across multiple compound networks, indicating their broad regulatory influence in adipocyte metabolism. In the capsaicin network (**Fig. 5a, Tab S3**), key hub genes included FASN, HIF1A, ESR1, STAT3, and MTOR, implicating pathways related to lipogenesis, adipocyte function, and metabolic signalling. The caffeic acid network (**Fig. 5b, Tab S4**) revealed hub genes such as STAT3, ESR1, NFE2L2, and NOS2, pointing to roles in oxidative stress, inflammation, and lipolysis. Additional nodes like PTGS2, PTPN1, and JAK2 underscore a possible anti-inflammatory contribution to its mode of action. In the camphor network **(Fig. 5c, Tab S2**), prominent hubs included PPARA, FABP4, NOS2, and PPARG, reflecting direct involvement in lipid metabolism and adipogenic regulation. The presence of VDR, P2RX4, and NR1H4 suggests a potential role in thermogenesis and nuclear receptor-mediated signalling.

**Figure 5.**
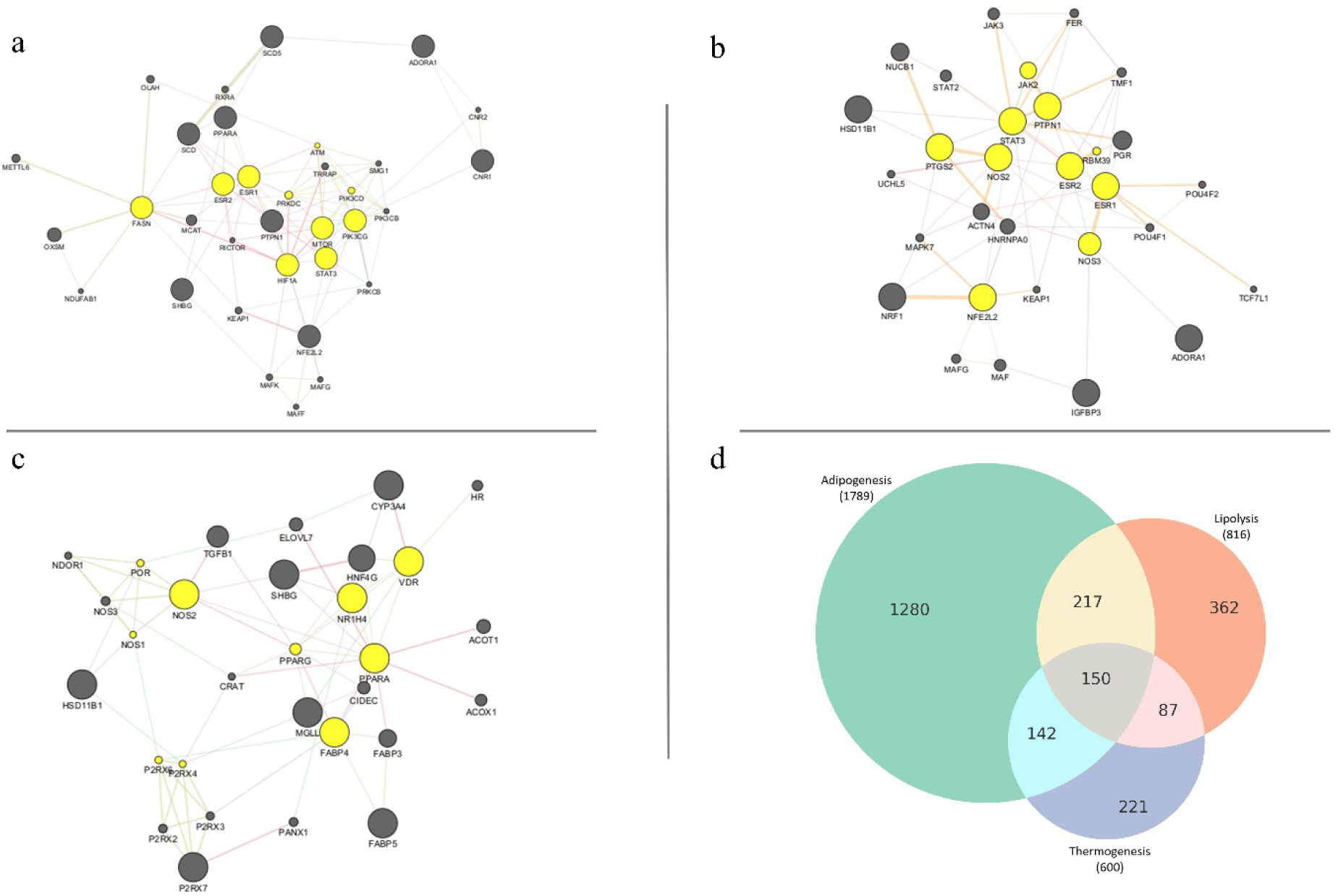
Network pharmacology analysis identifies hub genes targeted by capsaicin, camphor, and caffeic acid in adipocyte metabolic regulation. **(a – c)** Protein – protein interaction (PPI) networks for capsaicin **(a)**, caffeic acid **(b)**, and camphor **(c)** were constructed from overlapping predicted compound targets and genes associated with adipogenesis, lipolysis, and thermogenesis. Key shared regulators such as STAT3, ESR1/2, MTOR, and PPARG were consistently ranked across compounds, indicating central roles in metabolic signalling. **(d)** Venn diagram showing the overlap of genes involved in adipogenesis (n = 1789), lipolysis (n = 816), and thermogenesis (n = 600), with 150 genes common to all three processes. These genes formed the basis for compound target interaction and hub gene identification.

Overall, the integrated network analysis and functional mapping emphasize a core regulatory framework modulated by capsaicin, camphor, and caffeic acid. These findings provide mechanistic insight into how these bioactive compounds influence adipogenesis, lipolysis, and thermogenesis pathways. Graphical representations of the PPI networks for each compound (**Fig. 5a–c**) highlight the top hub genes, illustrating the interaction landscapes underpinning their metabolic actions.

## 4. Discussion

Adipogenesis is a highly regulated biological process involving a cascade of transcriptional and metabolic process that govern the conversion of preadipocytes into mature lipid storing adipocytes. This transition is tightly controlled by key transcription factors such as PPARγ and C/EBPα, which orchestrate the expression of genes responsible for lipid accumulation, insulin sensitivity, and energy homeostasis^22, 23^. Disruption of adipogenic signalling is implicated in metabolic disorders including obesity, insulin resistance, and type 2 diabetes^24^. In the present study, we evaluated the anti adipogenic and lipolytic potential of capsaicin, camphor, and caffeic acid as well as their combination in human SGBS adipocytes using an integrative approach combining *in vitro* assays, gene expression, and network pharmacology.

Adipogenesis assay revealed that all three compounds significantly inhibited lipid accumulation compared to vehicle controls. These findings are consistent with qPCR data, where key lipogenic transcription factors such as PPARγ and C/EBPα are downregulated following treatment. Such anti-adipogenic effects have previously been attributed to capsaicin’s and caffeic acid ability to modulate transcriptional networks and ERK/AMPK pathway respectively^25, 26^. Our study uniquely demonstrates a synergistic interaction between capsaicin and caffeic acid, providing new insight into their combination approach for more effective inhibition of adipogenesis. Similarly, lipolysis assays demonstrated increased glycerol release across all treatment groups. In support, qPCR showed significant upregulation of HSL across all treatment groups, most notably with capsaicin, supporting its direct effect on releasing stored fatty acids. These observations align with previous studies reporting capsaicin induced lipolysis through activation of adrenergic signalling and intracellular calcium elevation in iWAT^27^. Notably, TRPV1, molecular target of capsaicin, emerged as a common mediator in both adipogenic and lipolytic contexts. This agrees with previous findings showing that TRPV1 activation enhances energy expenditure and stimulates UCP1 mediated thermogenesis in adipose tissue^28, 29^. Interestingly, our results demonstrate that both camphor and caffeic acid also induce TRPV1 expression, suggesting possible indirect activation pathways or sensitization of TRP channels, further contributing to their observed metabolic effects^30, 31^. Our *in silico* network pharmacology approach revealed both overlapping and distinct hub genes associated with each compound. Genes such as STAT3, MTOR, and ESR1 were common across all three compounds, reflecting a shared axis of transcriptional control and metabolic regulation. These genes are known to mediate processes including inflammation, cell survival, and metabolic reprogramming hallmarks of adipose tissue remodelling^32, 33^. Capsaicin uniquely targeted genes such as FASN, HIF1A, and PIK3CD all of which are central regulators of lipid storage, adipocyte differentiation, and thermogenesis. VarElect analysis further substantiated the relevance of these hubs by directly linking them to lipolysis, adipogenesis, and thermogenesis with high phenotype matching scores. Conversely, camphor’s network was uniquely enriched for hubs which are the key markers of adipogenesis such as PPARγ, PPARα, and FABP4, highlighting pathways related to hormonal signalling, adipokine regulation, and metabolic control. These insightful observations are in partial agreement with camphor’s reported ability to influence MAPK and Wnt/β-catenin pathways^34^ and suggest a role in promoting thermogenesis and modulating adipocyte function. Caffeic acid’s regulatory network prominently featured STAT3, ESR1, NOS2, and PTGS2 indicating involvement in hypoxia responsive signalling, nitric oxide metabolism, and inflammatory regulation. These findings suggest that caffeic acid may exert a dual role in modulating adipocyte metabolism and resolving low grade inflammation, a feature with therapeutic potential in obesity associated metabolic dysfunction^35^.

Collectively, these findings indicate that capsaicin, camphor, and caffeic acid inhibit adipogenesis and enhance lipolysis via a multifaceted mechanism involving transcriptional repression, TRPV1 signalling, and modulation of key metabolic regulators. The consistency between experimental data and network based predictions underscore the translational value of these compounds as candidate agents for anti obesity therapies and its linked metabolic disorders e.g. diabetes. Our results demonstrated that the combination treatments which are largely unexplored effectively enhance lipolysis, suppress adipogenesis and alter gene expression level towards a metabolically active beige phenotype.

In support of these findings, some mechanistic aspects yet remain to be clarified. TRPV1 is reported to mediate key metabolic effects^36^, its specific contribution within the signalling network requires functional validation using genetic loss of function approaches, such as CRISPR/Cas9-mediated knockout or siRNA silencing. While downstream components like Ca²⁺-dependent kinases, AMPK, and UCP1 have been implicated in TRPV1 linked thermogenic signalling, the precise sequence and regulatory dynamics of this cascade remain incompletely characterized^37.^ Finally, to establish translational relevance, validation through *in vivo* studies is crucial, particularly using high fat diet induced obese mice, to assess whole body metabolic outcomes, compound bioavailability, and tissue specific effects.

## 5. Conclusion

Our findings demonstrate that capsaicin, camphor, and caffeic acid modulate adipocyte metabolism by inhibiting adipogenesis and promoting lipolysis in human SGBS cells. Gene expression analysis showed downregulation of key adipogenic genes including PPARG and CEBPA and upregulation of the key lipolytic marker HSL. TRPV1 activation, indicated by immunofluorescence, suggests its potential involvement. Importantly, combination treatments exhibited complementary effects on gene regulation and lipid metabolism, highlighting their potential for promoting metabolic remodelling in adipocytes.

## Acknowledgements

This work was supported under European Innovation Council project, BRAINSTORM, call HORIZON-EIC-2022-PATHFINDEROPEN-01, under Grant Agreement n. 101099355. We are grateful to Prof. Dr. Kristian Franze (FAU and MPZPM) and his lab, especially to Ms. Barbara Reischl, for access to their lab and assistance in performing qRT-PCRs.

## Author contribution

UA wrote the manuscript, designed the experiments, prepared the figures, and conducted the statistical analysis. DG provided intellectual guidance throughout the study, obtained funding and reviewed the manuscript. MW provided the SGBS cells and reviewed the manuscript.

## Data Availability

The materials and raw data are available from authors upon reasonable request.

